# No Evidence for Seasonal Variations in Fatigue, Sleepiness, and Insomnia Symptoms: Spring Fatigue is a Cultural Phenomenon rather than a Seasonal Syndrome

**DOI:** 10.1101/2025.09.27.678954

**Authors:** Christine Blume, Albrecht Vorster

## Abstract

Although not as prominent as in other animals, also humans experience seasonal variations in for example sleep duration and circadian processes. These variations are likely primarily driven by changes in photoperiod length. Anecdotally, a relevant number of people report experiencing fatigue and low energy levels particularly during spring – at least in Germany, Switzerland, and Austria. Thus, this phenomenon is commonly referred to as “spring fatigue”. However, scientific evidence for such a seasonal syndrome is largely missing.

We thus investigated temporal variations in fatigue, daytime sleepiness, insomnia symptoms, and sleep quality through an online survey including repeated (i.e., every six weeks) assessments of the same individuals over the course of one year. We hypothesised that fatigue and daytime sleepiness would be higher during shorter photoperiods. We further expected lower sleep quality and more severe insomnia symptoms under shorter photoperiods. Additionally, we explored variations with photoperiod change, across months, and seasons. Hypotheses were tested using Bayesian linear mixed-effects models. The study and analyses were pre-registered.

Between April 2024 and September 2025, 418 adults (80% women) completed at least two assessments. Nearly half of participants (47 %) reported experiencing spring fatigue. However, repeated assessments across one year showed no evidence for seasonal or monthly variations in fatigue, sleepiness, insomnia symptoms, or sleep quality. Fatigue during day-to-day activities decreased with longer photoperiods but was independent of photoperiod change.

Overall, the results provide evidence against spring fatigue as a genuine seasonal phenomenon. The discrepancy between high self-reports of the phenomenon and stable longitudinal patterns suggests that spring fatigue may reflect cultural labelling and result from cognitive-perceptual biases rather than reflecting a genuine seasonal syndrome.

## Introduction

In Germany, Switzerland, and Austria, the term “spring fatigue” (German: “Frühjahrsmüdigkeit” or “Frühlingsmüdigkeit”) is commonly used to describe the fatigue or lack of energy that many individuals report during spring. In a representative survey conducted by the Emnid institute that is often cited in this context, 22% of men and 39% of women reported suffering from “spring fatigue” (Kapp, 2022). However, it remains unclear whether fatigue is indeed stronger during spring compared to other seasons and, more generally, to what extent fatigue levels vary with photoperiod length, a plausible modulating factor.

Although seasonal variations in biological rhythms and sleep are less pronounced in humans than in animals, they are nonetheless present. Under natural lighting conditions (here: one week of camping in the Rocky Mountains), the internal biological night as measured by the duration of melatonin secretion was found to last about 4.4 hours longer than in summer. This effect was mainly due to the extension of the biological night into the morning hours and the effect was mainly attributed to the short photoperiod during winter (Stothard et al., 2017). Under electric lighting conditions, the authors of this study did not find seasonal differences in the duration of the biological night, suggesting that seasonal effects may likely be less pronounced in modern societies. In terms of sleep, Yetish and colleagues reported that in preindustrial societies in Tanzania, Namibia, and Bolivia sleep lasts about 1 hour longer in winter than in summer (Yetish et al., 2015). However, also in modern societies, seasonal variations in sleep have been reported. For example, a Swedish cohort study reported that participants interviewed in summer (i.e., June-August) were more likely to report ≤ 6 hours of sleep than participants surveyed in autumn (i.e., September-November; Titova et al., 2022). A large study involving 1856 participants aged 20-79 from Japan by Hashizaki et al. (2018) used contactless biomotion sensors to obtain objective sleep data across three years. While sleep onset remained largely consistent across seasons, sleep offset occurred later in winter an effect that was most pronounced on free days, when sleep times are not constrained by work obligations. These findings align well with delayed melatonin offset reported by Stothard et al. (2017). The finding that people tend to sleep more during the winter months was confirmed in another prospective longitudinal study from Japan by Suzuki et al. (2019), who found sleep to last about 11 minutes longer in winter than in summer. Similar effects were observed in the US, where sleep duration decreases particularly during spring compared to winter (Mattingly et al., 2021) and subtle seasonal variations in sleep architecture have been reported in neuropsychiatric patients (Seidler et al., 2023).

Findings regarding seasonal variations in the prevalence of sleep problems are less consistent. In the Swedish cohort study, complaints about sleep problems (i.e., falling asleep and disturbed sleep) were more common among those interviewed during spring (i.e., March-May) than during autumn (Titova et al., 2022). Furthermore, in the study by Hashizaki and colleagues sleep quality – measured by wakefulness after sleep onset and sleep efficiency (i.e., the duration of sleep compared to the time spent in bed) – was lowest during mid-summer (Hashizaki et al., 2018). Conversely, findings from the UK indicated that the time spent in bed was longer during winter while self-reported sleep times did not vary across seasons, which effectively could result in decreased sleep efficiency during the winter (O’Connell et al., 2014). A study from Norway also reported that insomnia symptoms (i.e., sleep onset problems and daytime impairments) were more common during winter compared to summer (Pallesen et al., 2001), which is in line with a comprehensive population health survey conducted in the region of Tromsø north of the arctic circle (FRIBORG et al., 2012).

However, none of these studies investigated seasonal variations in fatigue or daytime sleepiness. In a sample of individuals with rheumatoid arthritis, Feldthusen and colleagues reported that fluctuations in fatigue were stronger during winter (Feldthusen et al., 2016). However, in patients suffering from multiple sclerosis, fatigue was stronger during summer, possibly related to higher outdoor temperatures (Grothe et al., 2022). The only study conducted with healthy individuals reported increased fatigue, depressive mood, and problems falling asleep during winter in a sample from northern Norway (69° latitude), but not from Ghana (5° latitude; FRIBORG et al., 2012).

Here, we therefore aimed to systematically investigate seasonal variations in fatigue, insomnia symptoms, daytime sleepiness, and sleep quality with an online survey with up to 9 assessments across a whole year (i.e., repeated assessments every 6 weeks). In line with previous research, we hypothesized that period length would be the main modulator and more specifically that fatigue levels and daytime sleepiness would be higher, sleep quality lower and insomnia symptoms stronger when photoperiod is shorter.

## Methods

### Preregistration

The project including hypotheses and the analytic strategy were preregistered on Open Science Framework (OSF) on 19 March 2025 under the following link: osf.io/f7sxh.

### Sample

The final sample included 418 participants with a median of 8 completed assessments. Of the participants, 80 % were women, 14 % men, 4 % diverse and 1 % preferred not to disclose. The median age was 32 years (range: 18-87 years, interquartile range 25-43 years). Participants came from Germany (58 %), Switzerland (32 %), and Austria (10 %). They were recruited through the investigators’ networks and outreach activities on public radio and LinkedIn as well as an advertisement on Instagram. Participants had to be at least 18 years old and provide informed consent. The study protocol was approved by the cantonal ethics committee (Ethikkommission Nordwest- und Zentralschweiz; 2024-00478).

### Data acquisition

The study was an online questionnaire, which was made available using the REDCap software (Harris et al., 2019; Harris et al., 2009). It was open between April 4, 2024, and September 23, 2025. Following registration, participants were automatically invited to answer the questionnaires every 6 weeks for the duration of one year, i.e. 9 times in total to obtain good coverage across all seasons. In case participants skipped one assessment, they were still automatically invited for the next assessment. At the first assessment, they answered questions regarding demographics, socioeconomic status, country/canton of residence, job status (e.g., percentage of the contract), shift work, whether they have chronic illnesses, etc. During follow-up visits, they were asked whether there had been any changes regarding these topics, e.g. whether their weekly workload changed or whether they had started working in shifts. During the first as well as all eight follow-up assessments, participants were moreover invited to answer established questionnaires to evaluate primary endpoints were self-reported fatigue, daytime sleepiness, and self-reported sleep quality. At each assessment, participants were asked to rate their symptoms during the past four weeks. Self-reported fatigue was assessed with the “Fatigue Severity Scale” (FSS; Valko et al., 2008). This is a nine-item scale asking participants to rate their agreement to statements regarding fatigue during the past four weeks. A mean FSS score >4 is indicative of meaningful fatigue/daytime tiredness. Additionally, we assessed fatigue with a single question asking how fatigued participants generally felt during everyday activities during the preceding four weeks. Daytime sleepiness was assessed with the “Epworth Sleepiness Scale” (ESS; Bloch et al., 1999). In this eight-item scale, participants rate how likely it was that they would fall asleep during certain activities during the past four weeks (e.g., reading, when driving a car and stopping because of a traffic jam for several minutes). Subjective sleep quality was assessed with the “Bernese Sleep Health Questionnaire” (Vorster et al., 2024). In this questionnaire, participants rate a list of 28 symptoms relating to various sleep disorders regarding the frequency with which they occurred during the past three months. Importantly, as participants will answer the questionnaire every six weeks, we will ask to what extent the symptoms occurred during the past four weeks. However, this should impact the measurement of subjective sleep quality beyond the time frame for which conclusions can be drawn. The questionnaire includes the (GAD-2; Kroenke et al., 2007), a 2-item anxiety scale, and a 2-item depression scale (PHQ-2; Kroenke et al., 2003). Additionally, we included two questions regarding room temperature as this may affect seasonal variations in sleep, which however were not included in the sum score. Finally, insomnia symptoms were assessed with the “Insomnia Severity Index” (Dieck et al., 2018). In this questionnaire, participants rate possible insomnia symptoms regarding their severity during the past four weeks. Note that usually, the questionnaire asks for symptoms during the past two weeks. However, extending the period to four should not affect validity as the minimum duration of symptoms for an insomnia diagnosis according to international classifications of sleep disorders is anyway four weeks. To assess important control variables such as chronotype, social jetlag, light exposure, participants also filled in the Munich Chronotype Questionnaire (Ghotbi et al., 2019; Roenneberg et al., 2003). For a full code book, please see the preregistration on OSF. Additionally, the authors are happy to help with a translation as the code book is in German (as was the survey).

### Data reduction and statistical analysis

The data were downloaded from the REDCap and further processed in R version 4.4.2 (R Core Team, 2024).

We first removed datasets, where the first assessment was not completed or which did not include at least two completed assessments in total. The relevant values for each questionnaire (i.e., FSS, ISI, ESS, BSHQ, MCTQ) were computed in line with the instructions for each questionnaire. For the MCTQ, we corrected invalid data, e.g. where participants had entered 11:00 instead of 23:00 for their bedtime. Additionally, we corrected the “times spent under the open sky”, where people for instance entered “40min” instead of just “40” although they had been instructed to only enter the number in minutes. If they had entered a range of numbers, we took the mean value. For values smaller than 5, indicating they spend less than 5 minutes under the open sky on average, we assumed that people had erroneously entered a number for hours spent under the open sky and corrected this accordingly. The information on the photoperiod was calculated based on the federal state or canton people lived in. Specifically, we used the latitude of the geographical centre of the state or canton and calculated the photoperiod length for the date when the questionnaire had been filled in using the ‘photoperiod’ function from the ‘meteor’ package (Hijmans, 2023). For the ‘season’ variable, assessments were grouped as follows: March–May as spring, June–August as summer, September–November as autumn, and December–February as winter.

We analysed the data for the confirmatory and exploratory analyses using Bayesian linear mixed models as implemented in the ‘BayesFactor’ package (Morey & Rouder, 2019). Specifically, we compared the model including the effect of interest, for example photoperiod length, month, or season, against the model without the effect of interest. To test interactions, we compared the model including the interaction to the model with the additive effect. Additionally, the models included the factor “assessment number” (i.e., 1^st^-9^th^ assessment) to account for repeated assessments and participant ID and gender were entered as random factors. For all analyses, we report BF_10_, i.e., the likelihood of the data under H1 compared to H0, which results from the model comparison outlined above. The interpretation of the BF_10_ follows the suggestions by Jeffreys (1961), which we also summarise in the table below.

**Table 1:** Interpretation of Bayes Factors according to Jeffreys (1961).

| Bayes Factor | Interpretation |  |
| --- | --- | --- |
| **BF < 1/100** | **Extreme Evidence** | In favour of<br>H0 |
| **1/30 > BF ≥ 1/100** | **Very strong Evidence** |  |
| **1/10 > BF ≥ 1/30** | **Strong Evidence** |  |
| 1/3 > BF ≥ 1/10 | Moderate Evidence |  |
| **1 > BF ≥ 1/3** | Anecdotal Evidence |  |
| BF = 1 | *No Evidence* |  |
| **1 < BF ≤ 3** | Anecdotal Evidence | In favour of<br>H1 |
| 3 < BF ≤ 10 | Moderate Evidence |  |
| **10 < BF ≤ 30** | **Strong Evidence** |  |
| **30 < BF ≤ 100** | **Very strong Evidence** |  |
| **BF > 100** | **Extreme Evidence** |  |

Bayes factors in bold are deemed conclusive, all other evidence is inconclusive. Abbreviations: BF = Bayes Factor; H0 = null hypothesis; H1 = alternative hypothesis.

Following general recommendations (van Doorn et al., 2021), error percentages below 20% were deemed acceptable (default: 20 000 iterations) as this should result in the same qualitative conclusion.

## Results

Overall, 47 % of the participants reported suffering from spring fatigue.

### Fatigue Severity

The severity of reported fatigue as assessed with the Fatigue Severity Scale (FSS) was generally high with a mean fatigue level of 4.0 ± 1.4 across all assessments, where a score larger than 4 is deemed clinically relevant. The fatigue experienced during day-to-day activities during the preceding 4 weeks as assessed with a single question and a visual analogue scale ranging from “not at all fatigued” (corresponding to 0) to “extremely fatigued” (corresponding to 100) yielded a mean of 57.5 ± 24.4.

There was moderate-to-strong evidence against variations in fatigue severity as assessed with the FSS due to photoperiod length. The data was approximately 10 times more likely under the H0 than under the H1 (BF_10_ = 0.1), which is close to the threshold for strong evidence according to Jeffreys (1961). We also explored the relationship between variations in fatigue severity and the rate of change of the photoperiod length, which yielded strong evidence in favour of the H0, that is, no effect (BF_10_ = 0.09), the same was true for the interaction of photoperiod change and season (BF_10_ = 0.015; very strong evidence). Additionally, there was extreme evidence against variations across months (BF_10_ < 0.001) and very strong evidence against variations across seasons (BF_10_ = 0.02). Supplementary Tables 1–4 present the posterior mean intercepts and slopes for photoperiod length and photoperiod change as well as the posterior mean intercept and estimated deviations for months and seasons. Overall, exploratory analyses of the influence of chronotype yielded moderate to extreme evidence against an interaction of fatigue severity with chronotype varying with photoperiod lengths, across months, or seasons (photoperiod length: BF_10_ = 0.13; months: BF_10_ < 0.001; season: BF_10_ = 0.19). Figure 1 A-C provides an illustration of the results.

**Figure 1.**
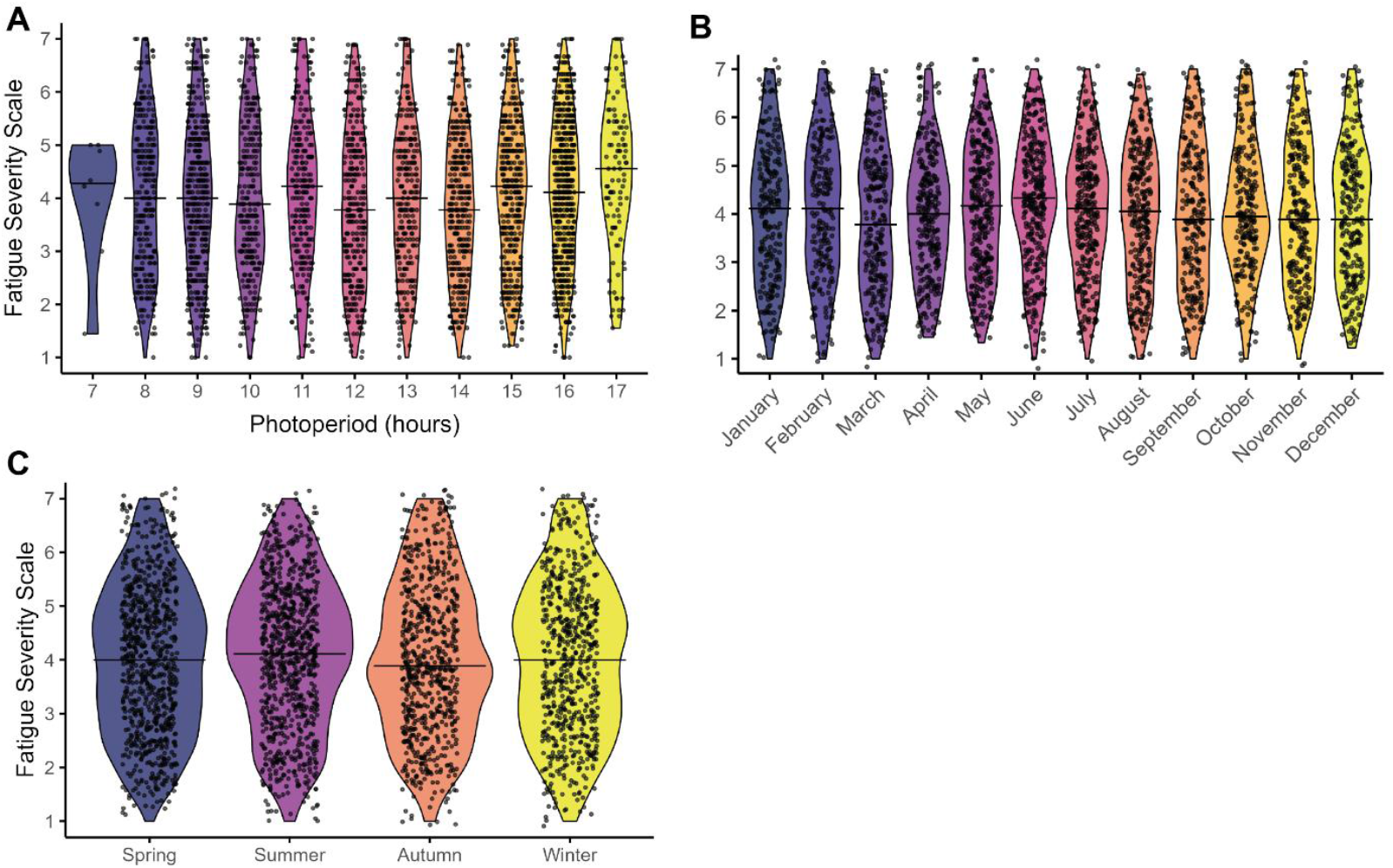
Variations in Fatigue Severity Scale scores across photoperiod lengths (A), months (B), and seasons (C). There was moderate-to strong evidence against variations with photoperiod length (BF = 0.1), extreme evidence against variations across months (BF < 0.001) and very strong evidence against variations across seasons (BF = 0.02). The width of each violin in the plots reflects the density of individual observations at each value. Individual participant data points are overlaid as dots. The solid horizontal line within each violin indicates the median score.

### Fatigue (single item, visual analogue scale)

The fatigue experienced during day-to-day activities during the preceding 4 weeks as assessed with a single question and a visual analogue scale (VAS) ranging from “not at all fatigued” (corresponding to 0) to “extremely fatigued” (corresponding to 100) yielded a mean of 57.5 ± 24.4. There was very strong evidence in favour of variations in fatigue experienced during day-to-day activities with photoperiod length (BF_10_ = 65.34) with fatigue decreasing with increasing photoperiod length. An exploratory analysis of the effect of rate of photoperiod change yielded strong evidence against such an effect (BF_10_ = 0.09) and the analysis of the interaction between photoperiod change and season yielded very strong evidence against such an effect (BF_10_= 0.02). Additionally, there was strong evidence against variations across months (BF_10_= 0.077) and moderate-to-strong evidence against variations across seasons (BF_10_ = 0.13). Supplementary Tables 5-8 present the posterior mean intercepts and slopes for photoperiod length and photoperiod change as well as the posterior mean intercept and estimated deviations for months and seasons. Figure 2 A-C provides an illustration of the results.

**Figure 2.**
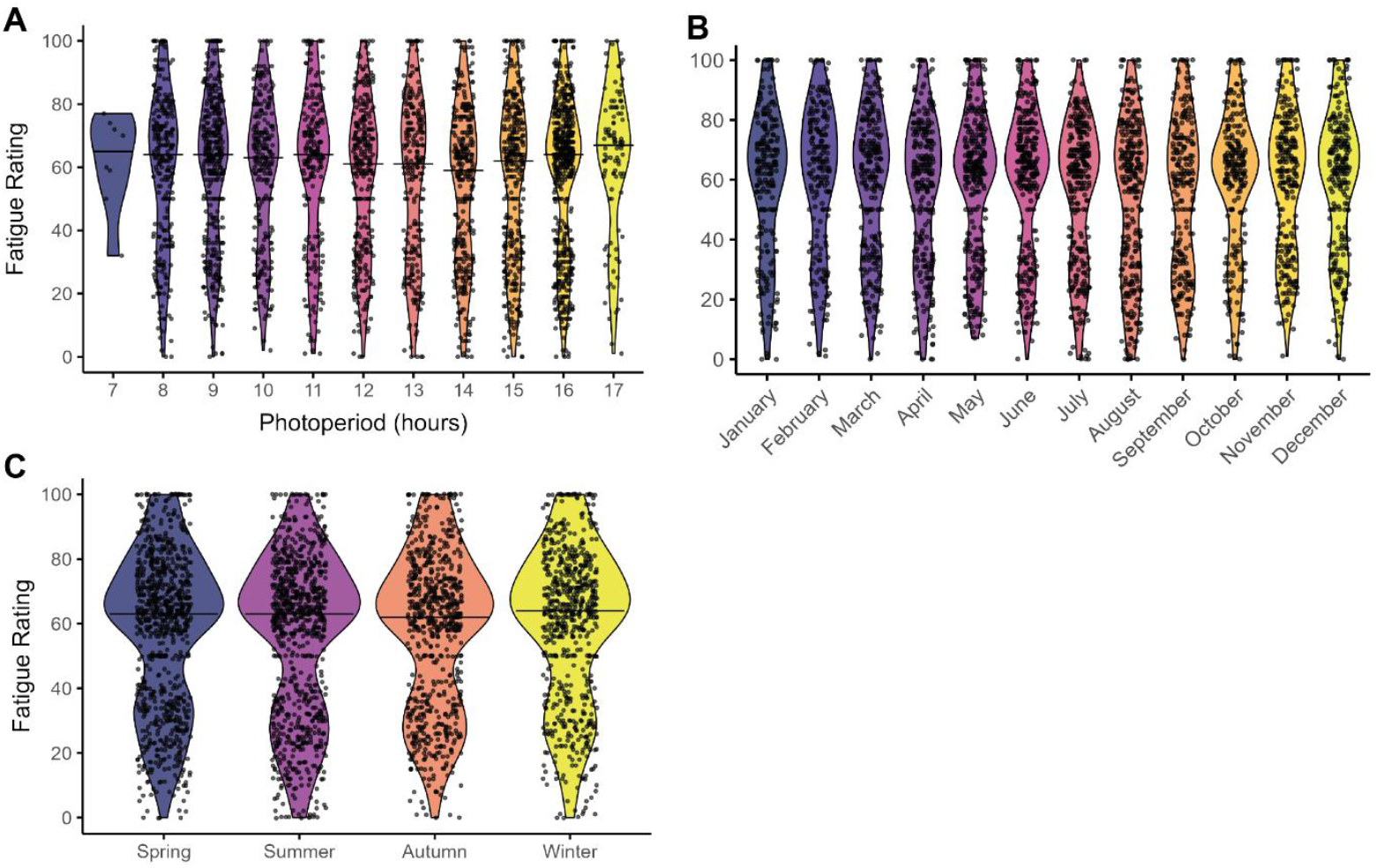
Variations in fatigue ratings during day-to-day activities on a visual analogue scale (VAS) across photoperiod lengths (A), months (B), and seasons (C). The VAS ranged from “not at all fatigued” (corresponding to 0) to “extremely fatigued” (corresponding to 100). There was very strong evidence in favour of variations with photoperiod length (BF = 65.34), strong evidence against variations across months (BF = 0.077) and moderate-to-strong evidence against variations across seasons (BF = 0.13). The width of each violin in the plots reflects the density of individual observations at each value. Individual participant data points are overlaid as dots. The solid horizontal line within each violin indicates the median score.

### Daytime Sleepiness

The average score on the Epworth Sleepiness Scale (ESS) assessing daytime sleepiness was 8.82 ± 4.63, where a value of 10 or larger is commonly accepted as a clinical cutoff. There was strong evidence against any effect of photoperiod length on daytime sleepiness: the data were approximately 13 times more likely under the null hypothesis than under the alternative (BF_10_ = 0.08). When examining the rate of change in photoperiod, results indicated moderate evidence for the null model (BF_10_ = 0.2). Moreover, there was extreme evidence against monthly variation (BF_10_ < 0.001) and strong evidence against seasonal variations (BF_10_ = 0.01). Supplementary Tables 9-13 provide an overview of the mean (intercept) and deviations from the intercept sampled from the posterior distribution for photoperiod length, months, and seasons.

### Insomnia Severity

The mean reported insomnia severity on the insomnia severity index (ISI) was 9.17 ± 5.52. This suggests that the average person in our sample had subthreshold insomnia, which corresponds to a sum score between 8 and 14 points.

Analyses provided strong evidence against an association between insomnia severity, as measured with the ISI, and photoperiod length. The data were 12.5 times more likely under the null hypothesis than under the alternative (BF_10_ = 0.8). When examining the rate of change in photoperiod, results indicated strong evidence for the null model, suggesting no effect (BF_10_ = 0.099). Moreover, we found extreme evidence against monthly variations (BF_10_ < 0.001) and strong evidence against seasonal variations (BF_10_ = 0.04). Supplementary Tables 13-16 report the posterior mean intercepts and slopes for photoperiod length and photoperiod change, along with posterior mean intercepts and estimated deviations for months and seasons.

### Sleep Health/Quality

On average, participants reported sleep health/quality values of 4.27 ± 1.51 on the Bernese Sleep Health Questionnaire (BSHQ). Values could range between 0 and 8 with higher values indicating better sleep health. Analyses yielded strong evidence against variations of BSHQ scores with photoperiod length, with the data being approx. 10 times more likely under the null model than under the alternative (BF_10_ = 0.098). Considering the rate of change in photoperiod, the results again supported the null hypothesis, pointing to no effect (BF_10_ = 0.07). In addition, the analyses revealed extreme evidence against differences across months (BF_10_ < 0.001) and seasonal effects (BF_10_ = 0.005). Supplementary Tables 17-20 present posterior mean intercepts and slopes for photoperiod length and change, as well as posterior mean intercepts and deviations for months and seasons.

### Exploratory Analyses

Beyond the confirmatory analyses, we explored variations in chronotype, sleep duration, and social jetlag across photoperiod lengths, months, and seasons. For these results, please see the Supplementary Material.

## Discussion

‘Frühjahrsmüdigkeit’ or ‘Frühlingsmüdigkeit’ (English: ‘spring fatigue’) is a widely used term, particularly in German-speaking countries Germany, Austria, and Switzerland. It refers to a perceived lack of energy and increased fatigue during day-to-day activities during spring. In our sample, nearly half of all participants (47 %) identified as being affected, underlining the phenomenon’s high subjective prevalence. Strikingly, however, the empirical findings collected here in regularly repeated assessments (i.e., every six weeks) across one year offer no support for such variations. Specifically, we found evidence for the absence of variations in fatigue levels during day-to-day activities and fatigue severity as a function of photoperiod length, calendar month, or season, with one exception: fatigue experienced during day-to-day activities decreased with longer photoperiods. Importantly, however, we found conclusive evidence against variations with photoperiod change – an effect that would have been expected if spring fatigue were a genuine phenomenon. Additionally, none of the investigated relationships were moderated by chronotype. Furthermore, we find overall conclusive evidence for the absence of temporal variations in daytime sleepiness, insomnia symptoms, or sleep health and quality. This stark contrast between subjective reports and objective data suggests that ‘spring fatigue’ does not reflect a genuine seasonal syndrome and rather points to the need for alternative explanatory frameworks.

We suggest that the high subjective prevalence could be the result of a ‘labelling effect’ (for an overview see Pohl, 2022) and further psychological biases such as an attribution and confirmation bias that the availability of a label can facilitate. More specifically, the term ‘Frühjahrsmüdigkeit’ is a culturally established, well known and frequently used label that can provide a readily available explanation for unspecific symptoms. It is known that labelling exerts robust effects on both memory and subjective experience, thereby enhancing recall and shaping perceptions. For instance, in an experiment by Bornstein (1976), a blue-green colour was remembered as bluer (or greener) when it was labelled as “blueish” (vs. “greenish”). Labels can also enhance the ability to reproduce a drawing (Hanawalt & Demarest, 1939). Additionally, wine was found to be judged as tasting better when it was labelled as more expensive (Plassmann et al., 2008). Likewise, a “FairTrade” label can improve the tastiness of a product (Lotz et al., 2013). In line with these findings, fatigue experienced during spring could be experienced as more intensely and remembered better than fatigue at other times, simply because there is a name – or label – for it. Furthermore, the existence of a label could contribute to an attribution and/or confirmation bias. Here, a familiar label allows to attribute unspecific symptoms of low energy to a seemingly obvious explanation: ‘spring fatigue’ (Kerstner et al., 2015). This again subjectively confirms the existence of the phenomenon. This attribution could, in turn, foster confirmation bias as it could lead people to pay attention to feelings of low energy particularly during spring, to ignore evidence challenging this interpretation, and to recall earlier episodes of ‘spring fatigue’ (Brewer & Treyens, 1981; Nickerson, 1998; Wason, 2021). The biases may further be increased by the high prevalence of the term ‘spring fatigue’ in the media and everyday conversations, which can prime individuals to direct their attention towards feelings of tiredness and interpret them in line with the primed concept or label, increasing reports of such symptoms particularly in spring (Kristjánsson & Ásgeirsson, 2019). Additionally, the wide media coverage creates a socially shared narrative that can be used to normalise the experienced symptoms. In fact, one of the reasons that led the authors to conduct this study were the recurring media inquiries about spring fatigue particularly in March and April. Thus, the label may enable collective understanding and possibly empathy, even if the causes underlying the experienced symptoms are diverse and not season-specific. Finally, ‘spring fatigue’ might be a culturally accepted way to reconcile the mismatch between feeling tired despite longer days and improving weather and thus help to reduce cognitive dissonance (Festinger, 1957; Harmon-Jones & Mills, 2019). During other seasons, other explanations such as summer heat possibly even affecting sleep or rainier days and less daylight in autumn and winter may suffice as explanations.

Nevertheless, there are other possible explanations for our findings. For example, it could be that ‘spring fatigue’ is a rather short-lived phenomenon that is not caught by asking people every six weeks about symptoms during the preceding four weeks. In this case, we may have missed the effect in our study design. While there is no data on how long symptoms of ‘spring fatigue’ usually last, the authors, who grew up in Germany and have lived in Switzerland and/or Austria for several years would intuitively expect symptoms to be sustained for some time. This aligns with the 2-4 weeks that are mentioned in a recent article in the magazine National Geographic (Kapp, 2022). Thus, there should have been a relevant effect on answers referring to the preceding four weeks. Further, we argue that even if symptoms were rather short-lived, none of the measured endpoints showed evidence of variation over time.

Our results do not exclude that specific subgroups such as hay fever sufferers (Bernstein et al., 2024; Santos et al., 2006) or individuals with depleted vitamin D levels at the end of winter (Nowak et al., 2016) experience spring-related changes in fatigue levels. Furthermore, some individuals may experience relevant fatigue during the days following the start of daylight saving time in Europe, which takes place on the last weekend in March as the clock change often acutely curtails sleep (Blume et al., 2019; Shapland et al., 2024). While such effects may well be diluted in the overall analysis, they however still do not provide a valid explanation for the discrepancy we found: the large number of individuals (47 %) who identified as being affected by ‘spring fatigue’ and the absence of such a pattern in the data collected every six weeks across a whole year. Focussing on possible explanations for this stark contrast also circumvents a potential limitation of the study: the non-representative sample, which was rather young, and included far more women than men.

In summary, while ‘spring fatigue’ is a common self-identified phenomenon in Germany, Austria, and Switzerland, our data provide evidence against systematic temporal variations in fatigue, daytime sleepiness, insomnia symptoms, or sleep quality across photoperiod lengths, months, or seasons. The contrast between high subjective prevalence and the absence of corresponding stable longitudinal patterns suggests that ‘spring fatigue’ is unlikely to represent a genuine seasonal syndrome. Instead, our findings point to the likely influence of a culturally established label, attribution and confirmation biases, effects of priming, and cognitive dissonance reduction in shaping symptom perception and recall. Future research should aim to disentangle these mechanisms, for example by experimentally testing how labels like ‘spring fatigue’ affect symptom reporting and recall in societies, where it is not a culturally established label. In addition, studies could identify subgroups that may be more susceptible to such effects (e.g. individuals with high health anxiety or greater sensitivity to media messaging). Finally, longitudinal designs with higher temporal resolution could help determine whether brief, transient increases in fatigue occur that were not captured by our assessment intervals.

## Funding

CB was funded by an Ambizione grant from the Swiss National Science Foundation awarded to her (Project number 201742). Additionally, the project was funded by a project grant from the German Society for Sleep Research and Sleep Medicine (DGSM) awarded to her.

## Author Contributions

Conceptualization: CB, supported by AV; Methodology: CB; Formal analysis: CB; Investigation: CB; Data curation: CB; Writing – original draft: CB; Writing – review & editing: AV; Visualization: CB; Project administration: CB; Funding acquisition: CB.

## Conflicts of Interest

C.B. has had the following commercial interests related to sleep and/or light: honoraria for invited talks and workshops from F.A. Hoffmann-La Roche AG, L’Oréal, Swissline Cosmetics, Ruby Hotels, and Vattenfall. C.B. is an elected member of the Daylight Academy.

## Notes

### Competing Interest Statement

The authors have declared no competing interest.

### Summary of Updates

Minor updates to the abstract and included author contributions statement.

