## Supplementary Material for "No Evidence for Seasonal Variations in Fatigue, Sleepiness, and Insomnia Symptoms: Spring Fatigue is a Cultural Phenomenon rather than a Seasonal Syndrome"

Beyond the confirmatory analyses, we explored variations in chronotype, sleep duration, and social jetlag across photoperiod lengths, months, and seasons.

*Sleep Duration*

Average sleep duration on workdays was 7.4 ± 1.17 hours. Analyses yielded strong evidence in favour of a variation of sleep duration on workdays with photoperiod length ($BF_{10}$ = 18.79) with sleep duration being shorter when photoperiod was longer. There was anecdotal evidence against variations across months ($BF_{10}$ = 0.37) and anecdotal evidence in favour of seasonal variations ($BF_{10}$ = 1.6). Supplementary Tables 21-23 present posterior mean intercepts and slopes for photoperiod length and change, as well as posterior mean intercepts and deviations for months and seasons.

Mean sleep duration on free days was 8.23 ± 1.27 hours. Analyses provided extreme evidence in favour of variations of free day sleep duration with photoperiod length ($BF_{10} > 1000$) with shorter sleep durations when photoperiod was longer. Furthermore, the results showed moderate evidence against variations across months ($BF_{10}$ = 0.31) and very strong evidence in favour of variations of free day sleep duration across seasons ($BF_{10}$ = 76.53) with shorter sleep durations during spring and summer. Supplementary Tables 24–26 report the posterior mean intercepts and slopes for photoperiod length and its rate of change, along with posterior mean intercepts and estimated deviations for months and seasons.

*Social Jetlag*

On average, participants reported social jetlag (SJL) derived from the MCTQ of 11.73 ± 17.75 minutes. Analyses yielded anecdotal evidence against variations of SJL across photoperiod length ($BF_{10}$ = 0.37). In addition, the analyses revealed extreme evidence against differences across months ($BF_{10} < 0.001$) and extreme evidence against seasonal effects ($BF_{10}$

= 0.009). Supplementary Tables 25-27 present posterior mean intercepts and slopes for photoperiod length and change, as well as posterior mean intercepts and deviations for months and seasons.

*Chronotype*

Average mid sleep on free days corrected for oversleep as assessed with the MCTQ was 03:23 ± 1.86 hours. Analyses yielded moderate evidence against variations of chronotype with photoperiod length ($BF_{10}$ = 0.12). There was extreme evidence against variations across months ($BF_{10}$ = 3.52) and seasonal variations ($BF_{10}$ = 0.003). Supplementary Tables 28-30 present posterior mean intercepts and slopes for photoperiod length and change, as well as posterior mean intercepts and deviations for months and seasons.

### Supplementary Tables

#### *Fatigue Severity*

**Suppl. Table 1:** Posterior mean intercept and slope for the effect of photoperiod length on the Fatigue Severity Scale (FSS). The intercept represents the expected outcome at the mean photoperiod length. The photoperiod slope quantifies the average change in the outcome per hour increase in daylight length.

| Effect | Estimate | SD | 95% CI (lower; upper) |
|---|---|---|---|
| Photoperiod length mean (intercept) | 4.3 | 0.43 | 3.47; 5.19 |
| Photoperiod length (slope) | 0.008 | 0.008 | -0.01; 0.02 |

Abbreviations: SD = Standard deviation of the estimate; CI = Credible Interval. For each condition, we report differences from the intercept.

**Suppl. Table 2:** Posterior mean intercept and slope for the effect of photoperiod change on the Fatigue Severity Scale (FSS). The intercept represents the expected outcome at the mean photoperiod length. The photoperiod slope quantifies the average change in the outcome per hour increase in daylight length.

| Effect | Estimate | SD | 95% CI (lower; upper) |
|---|---|---|---|
| Photoperiod change mean (intercept) | 4.3 | 0.45 | 3.43; 5.18 |
| Photoperiod change (slope) | -0.41 | 0.48 | -1.36; 0.54 |

Abbreviations: SD = Standard deviation of the estimate; CI = Credible Interval. For each condition, we report differences from the intercept.

**Suppl. Table 3.** Posterior mean intercept and estimated deviations from the intercept across months on the Fatigue Severity Scale (FSS). The intercept corresponds to the overall mean across all categories. Coefficients for individual categories indicate their deviation from this mean.

| Effect | Estimate | SD | 95% CI (lower; upper) |
|---|---|---|---|
| Month mean (intercept) | 4.3 | 0.43 | 3.48; 5.19 |
| January | -0.6 | 0.05 | -0.16; 0.05 |
| February | 0.04 | 0.06 | -0.07; 0.15 |
| March | -0.01 | 0.05 | -0.12; 0.09 |
| April | -0.01 | 0.05 | -0.11; 0.09 |
| May | -0.05 | 0.05 | -0.15; 0.44 |
| June | 0.08 | 0.05 | -0.02; 0.17 |
| July | 0.05 | 0.05 | -0.05; 0.15 |
| August | 0.001 | 0.05 | -0.09; 0.1 |

| | | | |
|---|---|---|---|
| September | -0.05 | 0.05 | -0.15; 0.05 |
| October | 0.03 | 0.05 | -0.08; 0.13 |
| November | 0.04 | 0.05 | -0.06; 0.15 |
| December | -0.06 | 0.05 | -0.16; 0.04 |

Abbreviations: SD = Standard deviation of the estimate; CI = Credible Interval. For each condition, we report differences from the intercept.

**Suppl. Table 4:** Posterior mean intercept and estimated deviations from the intercept across seasons on the Fatigue Severity Scale (FSS). The intercept corresponds to the overall mean across all categories. Coefficients for individual categories indicate their deviation from this mean.

| **Effect** | **Estimate** | **SD** | **95% CI (lower; upper)** |
|---|---|---|---|
| Season mean (intercept) | 4.3 | 0.45 | 3.43; 5.2 |
| Spring | -0.43 | 0.33 | -0.11; 0.02 |
| Summer | 0.06 | 0.32 | -0.002; 0.12 |
| Autumn | 0.02 | 0.34 | -0.43; 0.92 |
| Winter | -0.04 | 0.33 | -0.11; 0.02 |

Abbreviations: SD = Standard deviation of the estimate; CI = Credible Interval. For each condition, we report differences from the intercept.

#### *Fatigue (Single item, visual analogue scale)*

**Suppl. Table 5:** Posterior mean intercept and slope for the effect of photoperiod length on fatigue ratings (single item; visual analogue scale). The intercept represents the expected outcome at the mean photoperiod length. The photoperiod slope quantifies the average change in the outcome per hour increase in daylight length.

| **Effect** | **Estimate** | **SD** | **95% CI (lower; upper)** |
|---|---|---|---|
| Photoperiod length mean (intercept) | 61.2 | 8.75 | 44.61; 78.38 |
| Photoperiod length (slope) | -0.76 | 0.21 | -1.17; -0.35 |

Abbreviations: SD = Standard deviation of the estimate; CI = Credible Interval. For each condition, we report differences from the intercept.

**Suppl. Table 6:** Posterior mean intercept and slope for the effect of photoperiod change on fatigue ratings (single item; visual analogue scale). The intercept represents the expected outcome at the mean photoperiod length. The photoperiod slope quantifies the average change in the outcome per hour increase in daylight length.

| Effect | Estimate | SD | 95% CI (lower; upper) |
|---|---|---|---|
| Photoperiod change mean (intercept) | 61.46 | 8.8 | 44.25; 78.46 |
| Photoperiod change (slope) | 4.2 | 11.54 | -18.66; 26.42 |

Abbreviations: SD = Standard deviation of the estimate; CI = Credible Interval. For each condition, we report differences from the intercept.

**Suppl. Table 7.** Posterior mean intercept and estimated deviations from the intercept across months on fatigue ratings (single item; visual analogue scale). The intercept corresponds to the overall mean across all categories. Coefficients for individual categories indicate their deviation from this mean.

| Effect | Estimate | SD | 95% CI (lower; upper) |
|---|---|---|---|
| Month mean (intercept) | 61.67 | 8.57 | 44.88; 78.79 |
| January | 0.8 | 1.34 | -1.78; 3.45 |
| February | 2.93 | 1.39 | 0.29; 5.71 |
| March | 1.78 | 1.24 | -0.65; 4.23 |
| April | -2.0 | 1.26 | -4.44; 1.46 |
| May | -2.65 | 1.21 | -5.0; -0.31 |
| June | -0.35 | 1.19 | -2.7; 1.96 |
| July | -2.42 | 1.22 | -4.79; -0.03 |
| August | -3.18 | 1.21 | -5.55; -0.85 |
| September | -1.42 | 1.3 | -4.0; 1.12 |
| October | 1.72 | 1.3 | -0.8; 4.28 |
| November | 2.27 | 1.29 | -0.23; 4.79 |
| December | 2.49 | 1.32 | -0.03; 5.09 |

Abbreviations: SD = Standard deviation of the estimate; CI = Credible Interval. For each condition, we report differences from the intercept.

**Suppl. Table 8:** Posterior mean intercept and estimated deviations from the intercept across seasons on fatigue ratings (single item; visual analogue scale). The intercept corresponds to the overall mean across all categories. Coefficients for individual categories indicate their deviation from this mean.

| Effect | Estimate | SD | 95% CI (lower; upper) |
|---|---|---|---|
| Season mean (intercept) | 61.49 | 8.6 | 44.48; 78.87 |
| Spring | -1.36 | 0.8 | -2.94; 0.21 |
| Summer | -1.67 | 0.77 | -3.17; -0.14 |
| Autumn | 1.25 | 0.83 | -0.34; 2.93 |
| Winter | 1.78 | 0.84 | 0.11; 3.44 |

Abbreviations: SD = Standard deviation of the estimate; CI = Credible Interval. For each condition, we report differences from the intercept.

#### *Daytime Sleepiness*

**Suppl. Table 9:** Posterior mean intercept and slope for the effect of photoperiod length on daytime sleepiness as assessed with the Epworth Sleepiness Scale (ESS). The intercept represents the expected outcome at the mean photoperiod length. The photoperiod slope quantifies the average change in the outcome per hour increase in daylight length.

| **Effect** | **Estimate** | **SD** | **95% CI (lower; upper)** |
|---|---|---|---|
| Photoperiod length mean (intercept) | 7.36 | 1.26 | 4.93; 9.89 |
| Photoperiod length (slope) | 0.013 | 0.023 | -0.03; 0.06 |

Abbreviations: SD = Standard deviation of the estimate; CI = Credible Interval. For each condition, we report differences from the intercept.

**Suppl. Table 10:** Posterior mean intercept and slope for the effect of photoperiod change on daytime sleepiness as assessed with the Epworth Sleepiness Scale (ESS). The intercept represents the expected outcome at the mean photoperiod length. The photoperiod slope quantifies the average change in the outcome per hour increase in daylight length.

| **Effect** | **Estimate** | **SD** | **95% CI (lower; upper)** |
|---|---|---|---|
| Photoperiod change mean (intercept) | 7.38 | 1.3 | 4.83; 9.98 |
| Photoperiod change (slope) | -2.00 | 1.35 | -4.67; 0.64 |

Abbreviations: SD = Standard deviation of the estimate; CI = Credible Interval. For each condition, we report differences from the intercept.

**Suppl. Table 11.** Posterior mean intercept and estimated deviations from the intercept across months on daytime sleepiness as assessed with the Epworth Sleepiness Scale (ESS). The intercept corresponds to the overall mean across all categories. Coefficients for individual categories indicate their deviation from this mean.

| **Effect** | **Estimate** | **SD** | **95% CI (lower; upper)** |
|---|---|---|---|
| Month mean (intercept) | 7.38 | 1.22 | 4.97; 9.80 |
| January | -0.07 | 0.15 | -0.36; 0.22 |
| February | -0.02 | 0.16 | -0.34; 0.28 |

| | | | |
|---|---|---|---|
| March | -0.07 | 0.14 | -0.35; 0.21 |
| April | -0.26 | 0.15 | -0.55; 0.02 |
| May | -0.23 | 0.14 | -0.51; 0.04 |
| June | 0.26 | 0.14 | -0.01; 0.52 |
| July | 0.08 | 0.14 | -0.19; 0.35 |
| August | 0.12 | 0.14 | -0.14; 0.39 |
| September | 0.07 | 0.15 | -0.22; 0.35 |
| October | 0.03 | 0.15 | -0.27; 0.32 |
| November | 0.10 | 0.15 | -0.18; 0.40 |
| December | 0.01 | 0.15 | -0.28; 0.29 |

Abbreviations: SD = Standard deviation of the estimate; CI = Credible Interval. For each condition, we report differences from the intercept.

**Suppl. Table 12:** Posterior mean intercept and estimated deviations from the intercept across seasons on daytime sleepiness as assessed with the Epworth Sleepiness Scale (ESS). The intercept corresponds to the overall mean across all categories. Coefficients for individual categories indicate their deviation from this mean.

| **Effect** | **Estimate** | **SD** | **95% CI (lower; upper)** |
|---|---|---|---|
| Season mean (intercept) | 7.39 | 1.26 | 4.96; 9.95 |
| Spring | -0.24 | 0.10 | -0.43; -0.05 |
| Summer | 0.20 | 0.09 | 0.02; 0.38 |
| Autumn | 0.10 | 0.10 | -0.09; 0.29 |
| Winter | -0.06 | 0.10 | -0.25; 0.13 |

*Insomnia Severity*

**Suppl. Table 13:** Posterior mean intercept and slope for the effect of photoperiod length on insomnia severity as assessed with the Insomnia Severity Index (ISI). The intercept represents the expected outcome at the mean photoperiod length. The photoperiod slope quantifies the average change in the outcome per hour increase in daylight length.

| **Effect** | **Estimate** | **SD** | **95% CI (lower; upper)** |
|---|---|---|---|
| Photoperiod length mean (intercept) | 9.94 | 1.55 | 6.94; 13.08 |
| Photoperiod length (slope) | -0.18 | 0.03 | -0.81; 13.08 |

Abbreviations: SD = Standard deviation of the estimate; CI = Credible Interval. For each condition, we report differences from the intercept.

**Suppl. Table 14:** Posterior mean intercept and slope for the effect of photoperiod change on insomnia severity as assessed with the Insomnia Severity Index (ISI). The intercept represents the expected outcome at the mean photoperiod length. The photoperiod slope quantifies the average change in the outcome per hour increase in daylight length.

| **Effect** | **Estimate** | **SD** | **95% CI (lower; upper)** |
|---|---|---|---|
| Photoperiod change mean (intercept) | 9.93 | 1.5 | 7.05; 12.93 |
| Photoperiod change (slope) | -2.09 | 1.89 | -5.86; 1.52 |

Abbreviations: SD = Standard deviation of the estimate; CI = Credible Interval. For each condition, we report differences from the intercept.

**Suppl. Table 15.** Posterior mean intercept and estimated deviations from the intercept across months on insomnia severity as assessed with the Insomnia Severity Index (ISI). The intercept corresponds to the overall mean across all categories. Coefficients for individual categories indicate their deviation from this mean.

| **Effect** | **Estimate** | **SD** | **95% CI (lower; upper)** |
|---|---|---|---|
| Month mean (intercept) | 9.96 | 1.55 | 6.92; 13.01 |
| January | 0.14 | 0.21 | -0.27; 0.55 |
| February | 0.31 | 0.22 | -0.11; 0.74 |
| March | 0.19 | 0.20 | -0.59; 0.20 |
| April | 0.26 | 0.20 | -0.65; 0.14 |
| May | 0.32 | 0.19 | -0.70; 0.05 |
| June | 0.02 | 0.19 | -0.39; 0.34 |
| July | 0.04 | 0.19 | -0.41; 0.32 |
| August | 0.20 | 0.19 | -0.17; 0.58 |
| September | 0.09 | 0.20 | -0.31; 0.48 |
| October | 0.06 | 0.21 | -0.46; 0.36 |
| November | 0.11 | 0.20 | -0.28; 0.51 |
| December | 0.04 | 0.20 | -0.36; 0.43 |

Abbreviations: SD = Standard deviation of the estimate; CI = Credible Interval. For each condition, we report differences from the intercept.

**Suppl. Table 16:** Posterior mean intercept and estimated deviations from the intercept across seasons on insomnia severity as assessed with the Insomnia Severity Index (ISI). The intercept corresponds to the overall mean across all categories. Coefficients for individual categories indicate their deviation from this mean.

| Effect | Estimate | SD | 95% CI (lower; upper) |
|---|---|---|---|
| Season mean (intercept) | 9.96 | 1.52 | 7.01; 13.07 |
| Spring | -0.03 | 0.13 | -0.56; -0.66 |
| Summer | 0.06 | 0.12 | -0.18; 0.3 |
| Autumn | 0.07 | 0.13 | -0.19; 0.34 |
| Winter | 0.18 | 0.13 | -0.84; 0.44 |

#### *Sleep Health/ Quality*

**Suppl. Table 17:** Posterior mean intercept and slope for the effect of photoperiod length on sleep health/ quality as assessed with the Bernese Sleep Health Questionnaire (BSHQ; range 0-8 with higher values indicating better sleep). The intercept represents the expected outcome at the mean photoperiod length. The photoperiod slope quantifies the average change in the outcome per hour increase in daylight length.

| Effect | Estimate | SD | 95% CI (lower; upper) |
|---|---|---|---|
| Photoperiod length mean (intercept) | 3.89 | 0.51 | 2.88; 4.83 |
| Photoperiod length (slope) | 0.01 | 0.01 | -0.01; 0.03 |

Abbreviations: SD = Standard deviation of the estimate; CI = Credible Interval. For each condition, we report differences from the intercept.

**Suppl. Table 18:** Posterior mean intercept and slope for the effect of photoperiod change on sleep health/ quality as assessed with the Bernese Sleep Health Questionnaire (BSHQ; range 0-8 with higher values indicating better sleep). The intercept represents the expected outcome at the mean photoperiod length. The photoperiod slope quantifies the average change in the outcome per hour increase in daylight length.

| Effect | Estimate | SD | 95% CI (lower; upper) |
|---|---|---|---|
| Photoperiod change mean (intercept) | 3.89 | 0.49 | 2.91; 4.81 |
| Photoperiod change (slope) | 0.27 | 0.56 | -0.81; 1.33 |

Abbreviations: SD = Standard deviation of the estimate; CI = Credible Interval. For each condition, we report differences from the intercept.

**Suppl. Table 19.** Posterior mean intercept and estimated deviations from the intercept across months on sleep health/ quality as assessed with the Bernese Sleep Health Questionnaire (BSHQ; range 0-8 with higher values indicating better sleep). The intercept corresponds to the overall mean across all categories. Coefficients for individual categories indicate their deviation from this mean.

| **Effect** | **Estimate** | **SD** | **95% CI (lower; upper)** |
|---|---|---|---|
| Month mean (intercept) | 3.89 | 0.50 | 2.90; 4.82 |
| January | 0.04 | 0.06 | -0.08; 0.16 |
| February | -0.05 | 0.06 | -0.18; 0.07 |
| March | 0.00 | 0.06 | -0.11; 0.12 |
| April | 0.04 | 0.06 | -0.07; 0.16 |
| May | 0.05 | 0.06 | -0.06; 0.16 |
| June | -0.03 | 0.06 | -0.14; 0.08 |
| July | 0.03 | 0.06 | -0.08; 0.14 |
| August | 0.05 | 0.06 | -0.06; 0.16 |
| September | 0.01 | 0.06 | -0.11; 0.12 |
| October | -0.04 | 0.06 | -0.15; 0.09 |
| November | -0.08 | 0.06 | -0.20; 0.04 |
| December | -0.02 | 0.06 | -0.14; 0.10 |

Abbreviations: SD = Standard deviation of the estimate; CI = Credible Interval. For each condition, we report differences from the intercept.

**Suppl. Table 20:** Posterior mean intercept and estimated deviations from the intercept across seasons on sleep health/ quality as assessed with the Bernese Sleep Health Questionnaire (BSHQ; range 0-8 with higher values indicating better sleep). The intercept corresponds to the overall mean across all categories. Coefficients for individual categories indicate their deviation from this mean.

| **Effect** | **Estimate** | **SD** | **95% CI (lower; upper)** |
|---|---|---|---|
| Season mean (intercept) | 3.89 | 0.49 | 2.90; 4.84 |
| Spring | 0.05 | 0.04 | -0.03; 0.12 |
| Summer | 0.01 | 0.04 | -0.06; 0.08 |
| Autumn | -0.05 | 0.04 | -0.13; 0.02 |
| Winter | -0.00 | 0.04 | -0.08; 0.08 |

#### *Sleep duration on workdays*

**Suppl. Table 21:** Posterior mean intercept and slope for the effect of photoperiod length on sleep duration on workdays assessed with the Munich Chronotype Questionnaire. The intercept represents the expected outcome at the mean photoperiod length. The photoperiod slope quantifies the average change in the outcome per hour increase in daylight length.

| Effect | Estimate | SD | 95% CI (lower; upper) |
|---|---|---|---|
| Photoperiod length mean (intercept) | 7.31 | 0.36 | 6.60; 7.99 |
| Photoperiod length (slope) | -0.03 | 0.01 | -0.05; -0.01 |

Abbreviations: SD = Standard deviation of the estimate; CI = Credible Interval. For each condition, we report differences from the intercept.

**Suppl. Table 22.** Posterior mean intercept and estimated deviations from the intercept across months on sleep duration on workdays assessed with the Munich Chronotype Questionnaire. The intercept corresponds to the overall mean across all categories. Coefficients for individual categories indicate their deviation from this mean.

| Effect | Estimate | SD | 95% CI (lower; upper) |
|---|---|---|---|
| Month mean (intercept) | 7.31 | 0.36 | 6.61; 8.02 |
| January | 0.04 | 0.06 | -0.07; 0.15 |
| February | 0.08 | 0.06 | -0.03; 0.20 |
| March | 0.16 | 0.05 | 0.05; 0.26 |
| April | 0.08 | 0.05 | -0.03; 0.18 |
| May | -0.08 | 0.05 | -0.18; 0.03 |
| June | -0.10 | 0.05 | -0.20; 0.00 |
| July | -0.18 | 0.05 | -0.28; -0.07 |
| August | -0.08 | 0.05 | -0.18; 0.02 |
| September | 0.00 | 0.05 | -0.11; 0.11 |
| October | 0.04 | 0.06 | -0.07; 0.15 |
| November | 0.05 | 0.05 | -0.05; 0.16 |
| December | -0.02 | 0.05 | -0.13; 0.08 |

Abbreviations: SD = Standard deviation of the estimate; CI = Credible Interval. For each condition, we report differences from the intercept.

**Suppl. Table 23:** Posterior mean intercept and estimated deviations from the intercept across seasons on sleep duration on workdays assessed with the Munich Chronotype Questionnaire. The intercept corresponds to the overall mean across all categories. Coefficients for individual categories indicate their deviation from this mean.

| Effect | Estimate | SD | 95% CI (lower; upper) |
|---|---|---|---|
| Season mean (intercept) | 7.31 | 0.35 | 6.61; 7.98 |
| Spring | 0.05 | 0.03 | -0.01; 0.12 |
| Summer | -0.11 | 0.03 | -0.17; -0.04 |
| Autumn | 0.04 | 0.03 | -0.03; 0.11 |

| | | | |
|---|---|---|---|
| Winter | 0.02 | 0.03 | -0.05; 0.08 |

*Sleep duration on free days*

**Suppl. Table 24:** Posterior mean intercept and slope for the effect of photoperiod length on sleep duration on free days assessed with the Munich Chronotype Questionnaire. The intercept represents the expected outcome at the mean photoperiod length. The photoperiod slope quantifies the average change in the outcome per hour increase in daylight length.

| Effect | Estimate | SD | 95% CI (lower; upper) |
|---|---|---|---|
| Photoperiod length mean (intercept) | 8.08 | 0.38 | 7.32; 8.83 |
| Photoperiod length (slope) | -0.05 | 0.01 | -0.07; -0.03 |

Abbreviations: SD = Standard deviation of the estimate; CI = Credible Interval. For each condition, we report differences from the intercept.

**Suppl. Table 25.** Posterior mean intercept and estimated deviations from the intercept across months on sleep duration on free days assessed with the Munich Chronotype Questionnaire. The intercept corresponds to the overall mean across all categories. Coefficients for individual categories indicate their deviation from this mean.

| Effect | Estimate | SD | 95% CI (lower; upper) |
|---|---|---|---|
| Month mean (intercept) | 8.09 | 0.37 | 7.33; 8.80 |
| January | 0.08 | 0.06 | -0.04; 0.20 |
| February | 0.05 | 0.06 | -0.07; 0.18 |
| March | 0.07 | 0.06 | -0.05; 0.18 |
| April | -0.03 | 0.06 | -0.15; 0.08 |
| May | -0.14 | 0.06 | -0.26; -0.03 |
| June | -0.18 | 0.06 | -0.29; -0.07 |
| July | -0.15 | 0.06 | -0.26; -0.04 |
| August | -0.11 | 0.06 | -0.22; 0.00 |
| September | 0.09 | 0.06 | -0.03; 0.21 |
| October | 0.10 | 0.06 | -0.02; 0.22 |
| November | 0.12 | 0.06 | 0.00; 0.23 |
| December | 0.12 | 0.06 | -0.00; 0.24 |

Abbreviations: SD = Standard deviation of the estimate; CI = Credible Interval. For each condition, we report differences from the intercept.

**Suppl. Table 26:** Posterior mean intercept and estimated deviations from the intercept across seasons on sleep duration on free days assessed with the Munich Chronotype Questionnaire. The intercept corresponds to the overall mean across all categories. Coefficients for individual categories indicate their deviation from this mean.

| Effect | Estimate | SD | 95% CI (lower; upper) |
|---|---|---|---|
| Season mean (intercept) | 8.10 | 0.40 | 7.30; 8.85 |
| Spring | -0.04 | 0.04 | -0.11; 0.04 |
| Summer | -0.15 | 0.04 | -0.22; -0.08 |
| Autumn | 0.11 | 0.04 | 0.03; 0.18 |
| Winter | 0.08 | 0.04 | 0.002; 0.16 |

*Social Jetlag*

**Suppl. Table 25:** Posterior mean intercept and slope for the effect of photoperiod length on social jetlag assessed with the Munich Chronotype Questionnaire. The intercept represents the expected outcome at the mean photoperiod length. The photoperiod slope quantifies the average change in the outcome per hour increase in daylight length.

| Effect | Estimate | SD | 95% CI (lower; upper) |
|---|---|---|---|
| Photoperiod length mean (intercept) | 11.42 | 5.88 | -0.08; 23.19 |
| Photoperiod length (slope) | 0.30 | 0.16 | -0.02; 0.62 |

Abbreviations: SD = Standard deviation of the estimate; CI = Credible Interval. For each condition, we report differences from the intercept.

**Suppl. Table 26.** Posterior mean intercept and estimated deviations from the intercept across months on social jetlag assessed with the Munich Chronotype Questionnaire. The intercept corresponds to the overall mean across all categories. Coefficients for individual categories indicate their deviation from this mean.

| Effect | Estimate | SD | 95% CI (lower; upper) |
|---|---|---|---|
| Month mean (intercept) | 11.21 | 5.69 | -0.33; 22.61 |
| January | -1.55 | 1.07 | -3.70; 0.51 |
| February | 0.08 | 1.11 | -2.11; 2.22 |
| March | 0.88 | 1.02 | -1.14; 2.90 |
| April | -0.40 | 1.00 | -2.34; 1.55 |
| May | 2.02 | 0.98 | 0.12; 3.96 |
| June | 0.45 | 0.95 | -1.42; 2.33 |

| | | | |
|---|---|---|---|
| July | 0.57 | 0.98 | -1.38; 2.48 |
| August | 0.05 | 0.95 | -1.82; 1.91 |
| September | -0.85 | 1.03 | -2.87; 1.14 |
| October | -0.40 | 1.06 | -2.49; 1.68 |
| November | -0.10 | 1.03 | -2.15; 1.92 |
| December | -0.75 | 1.04 | -2.86; 1.26 |

Abbreviations: SD = Standard deviation of the estimate; CI = Credible Interval. For each condition, we report differences from the intercept.

**Suppl. Table 27:** Posterior mean intercept and estimated deviations from the intercept across seasons on sleep duration on social jetlag assessed with the Munich Chronotype Questionnaire. The intercept corresponds to the overall mean across all categories. Coefficients for individual categories indicate their deviation from this mean.

| **Effect** | **Estimate** | **SD** | **95% CI (lower; upper)** |
|---|---|---|---|
| Season mean (intercept) | 11.18 | 5.92 | -0.18; 22.90 |
| Spring | -0.53 | 0.67 | -1.85; 0.78 |
| Summer | 1.03 | 0.66 | -0.24; 2.32 |
| Autumn | 0.40 | 0.62 | -0.81; 1.60 |
| Winter | -0.90 | 0.68 | -2.23; 0.43 |

*Chronotype*

**Suppl. Table 28:** Posterior mean intercept and slope for the effect of photoperiod length on chronotype assessed with the Munich Chronotype Questionnaire. The intercept represents the expected outcome at the mean photoperiod length. The photoperiod slope quantifies the average change in the outcome per hour increase in daylight length.

| **Effect** | **Estimate** | **SD** | **95% CI (lower; upper)** |
|---|---|---|---|
| Photoperiod length mean (intercept) | 3.82 | 0.50 | 2.90; 4.85 |
| Photoperiod length (slope) | -0.01 | 0.01 | -0.03; 0.01 |

Abbreviations: SD = Standard deviation of the estimate; CI = Credible Interval. For each condition, we report differences from the intercept.

**Suppl. Table 29.** Posterior mean intercept and estimated deviations from the intercept across months on chronotype assessed with the Munich Chronotype Questionnaire. The intercept corresponds to the overall mean across all categories. Coefficients for individual categories indicate their deviation from this mean.

| Effect | Estimate | SD | 95% CI (lower; upper) |
|---|---|---|---|
| Month mean (intercept) | 3.82 | 0.49 | 2.91; 4.86 |
| January | 0.13 | 0.06 | 0.00; 0.26 |
| February | -0.00 | 0.07 | -0.13; 0.13 |
| March | -0.05 | 0.06 | -0.17; 0.07 |
| April | 0.01 | 0.06 | -0.11; 0.13 |
| May | 0.01 | 0.06 | -0.11; 0.12 |
| June | -0.03 | 0.06 | -0.14; 0.09 |
| July | -0.04 | 0.06 | -0.16; 0.07 |
| August | -0.03 | 0.06 | -0.14; 0.09 |
| September | -0.06 | 0.06 | -0.19; 0.06 |
| October | 0.06 | 0.06 | -0.07; 0.18 |
| November | 0.03 | 0.06 | -0.09; 0.15 |
| December | -0.02 | 0.06 | -0.15; 0.10 |

Abbreviations: SD = Standard deviation of the estimate; CI = Credible Interval. For each condition, we report differences from the intercept.

**Suppl. Table 30:** Posterior mean intercept and estimated deviations from the intercept across seasons on sleep duration on chronotype assessed with the Munich Chronotype Questionnaire. The intercept corresponds to the overall mean across all categories. Coefficients for individual categories indicate their deviation from this mean.

| Effect | Estimate | SD | 95% CI (lower; upper) |
|---|---|---|---|
| Season mean (intercept) | 3.82 | 0.50 | 2.92; 4.89 |
| Spring | -0.01 | 0.04 | -0.09; 0.06 |
| Summer | -0.04 | 0.04 | -0.11; 0.03 |
| Autumn | 0.01 | 0.04 | -0.07; 0.09 |
| Winter | 0.04 | 0.04 | -0.04; 0.12 |
